# Overhang Optimizer for Golden Gate Assembly (OOGGA)

**DOI:** 10.1101/2025.06.16.659877

**Authors:** S. Mukundan, M. S. Madhusudhan

## Abstract

Golden Gate cloning is a popular method for achieving ordered assembly of DNA fragments using complementary four-base pair overhangs. The success of this method relies on the ability of DNA ligases to ligate the different fragments. This ability is affected by the relative efficiency and accuracy with which the ligase recognizes and joins overhanging regions. In this study, we report a dynamic programming approach, called Overhang Optimizer for Golden Gate Assembly (OOGGA), that optimizes these overhangs for their accuracy and efficiency. OOGGA provides the theoretically optimal fragments for Golden Gate assembly, provided a DNA sequence, a length range of the fragments and/or the number of required fragments. As expected, OOGGA outperforms NEBridge SplitSet^®^ in both fidelity and efficiency metrics in a set of 3 example DNA sequences. OOGGA also provides support of degenerate nucleotide codes, options to exclude/include motifs for predictions and weights to bias the program for efficiency and fidelity.

## 1. Introduction

Golden Gate cloning is a molecular cloning method used to assemble multiple fragments of DNA in a specific orientation (Engler, Kandzia, and Marillonnet 2008). This is achieved using type IIs restriction enzymes such as BsaI and BsmBI. These restriction enzymes recognize specific 6 nucleotide sequence motifs, and cut the DNA at 1/5 sites downstream, creating a 4 base pair overhang at non-specific sites. These sites can be chosen such that DNA fragments share complementary overhangs that lead to assemblies facilitated by Watson-Crick base pairing. The nicks in these assemblies are ligated by DNA ligases (such as T4 and T7) present in the reaction to create progressively larger segments of DNA. This method is capable of accurately assembling more than 50 fragments of DNA (Pryor et al. 2020, 2022).

The accuracy of this method depends on sequence complementarity conferred by Watson-Crick base pairing. But there are exceptions to this rule that lead to mismatches, contributing to errors in assembly. Moreover, DNA ligases have sequence preferences that determine the efficiency and accuracy of ligation (Bilotti et al. 2022). The ligation efficiency for all 256 × 256 pairwise combinations of four base pair overhangs has been determined by single molecule sequencing (Potapov, Jennifer L Ong, et al. 2018; Potapov, Jennifer L. Ong, et al. 2018). Selecting high-ranking overhangs from these data (in terms of efficiency and fidelity) is necessary for robust experimental designs. Manual selection of these overhangs can get cumbersome and error-prone.

The tool, NEBridge SplitSet^®^ (SplitSet) uses data from Potatov et. al. to predict fragments of a DNA sequence with high-fidelity overhangs based on the number of required fragments. SplitSet’s Monte Carlo search approach identifies a high-fidelity set of overhangs from a large number of possible combinations (10^14^ for 10 overhangs) (Pryor et al. 2020). However, Monte Carlo is not deterministic and does not guarantee the best set of overhangs and the results from Monte Carlo simulations also differ from one run to another. Moreover, the method, while optimizing for fidelity, ignores the efficiency of ligation which is critical for applications like mutational screening, where the resulting larger populations are helpful and sometimes necessary.

Here we report a program called Overhang Optimizer for Golden Gate Assembly (OOGGA). This program uses dynamic programming to find the fragments of a given DNA sequence with the best set of overhangs, given the range of fragment length (the required number of fragments can also be provided). The efficiency and fidelity matrices, derived from the single molecule sequencing data generated by Potapov et al, are optimized in the process. We compare 5 fragments predicted for DNA sequences by both OOGGA and SplitSet. Compared to SplitSet, OOGGA predicts fragments with higher fidelity overhangs. Moreover, OOGGA can also optimize the efficiency of the overhangs.

## 2. Methods

### 2.1. Obtaining scores of efficiency and fidelity for overhangs

Metrics that quantify the efficiency and the fidelity of an overhang were calculated from the single molecule sequencing data generated by Potapov et al. The probability of an overhang undergoing ligation with its Watson-Crick counterpart (termed as a match; anything other than Watson-Crick ligations are termed mismatches) is considered as the metric for evaluating the fidelity (equation 1).

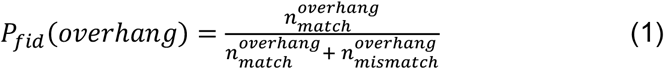

Here *n* is the number of reads obtained by Potapov et al. The value of the number of matching reads of an overhang divided by the value of the largest matching read among all the overhangs is considered as the metric of efficiency (equation 2). Note that this value ranges between 0-1 (equation 3).

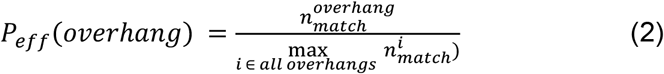

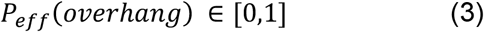

### 2.2 Overhang Optimizer for Golden Gate Assembly (OOGGA)

OOGGA employs dynamic programming to optimize overhang positions and the number of fragments for the Golden Gate assembly of a DNA sequence. The input to the program is the DNA sequence of interest and the size range of the fragments. The dynamic programming algorithm finds the combination of fragments that would maximize the probability of efficient and accurate assembly.

This is achieved by aligning overhang positions to the fragments while optimizing the net efficiency and fidelity of the assembly.

First, the number of fragments (*K*) is determined. If the number is not provided by the user, equation 4 is used to calculate the maximum possible number of fragments.

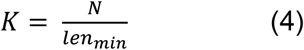

Where *N* is the number of base pairs in the DNA sequence and *K* is rounded up to the higher integer. *len*_*min*_ is the minimum length of the fragment (similarly *len*_*max*_ is the maximum length of the fragment). After this, four matrices with *K* columns and *N* rows are created, indexed by *i* and *j*, respectively. These are *S, P*_*eff*_, *P*_*fid*_ and *T* that store the total score, the e*ff*iciency score, the fidelity score, and the trace, respectively. The following recursive algorithm is used to populate these matrices.

### 2.3 Initiation

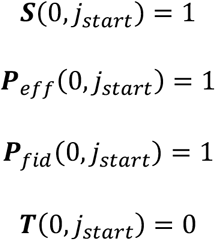

*j*_*start*_ is the position of the first fragment. This is by default set as 0. Assigning probabilities to multiple start sites can also optimise starting site fragments. Here *S, P*_*eff*_, *P*_*fid*_ and *T* are matrices with *K* columns and *N* rows. *S* stores the total score (see ‘Iteration’ section to see how score is computed) corresponding to the fragmentation at a given DNA position. *P*_*eff*_ and *P*_*fid*_ store the efficiency and fidelity that is used to compute the score. *T* stores the previous fragmentation site based on which the current fragmentation site is computed.

### 2.4 Iteration

Pseudocode for computing *S, P*_*eff*_, *P*_*fid*_ and *T*

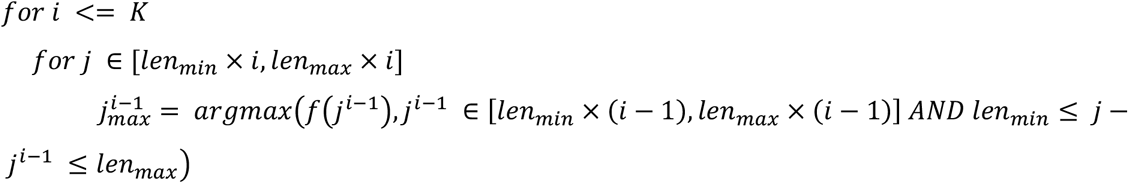

Where,

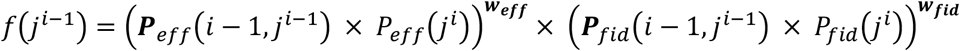

Using 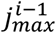 update the matrices

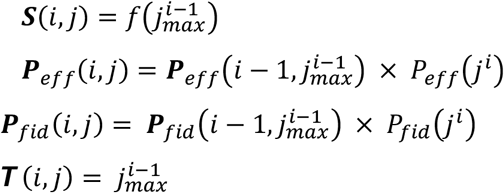

### 2.5 Termination

When *i > K*

Here, *w*_*eff*_ and *w*_*fid*_ are weights for efficiency and fidelity respectively.

### 2.6 Traceback

The top *n_trace* of the score of any overhang that can be part of the final fragment is selected as the starting point of the traceback.

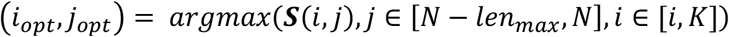

If the number of required fragments (*K*) is specified by the user, the following equation is used.

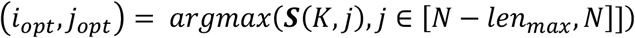

### 2.7 Time complexity

The number of total calculations is the following.

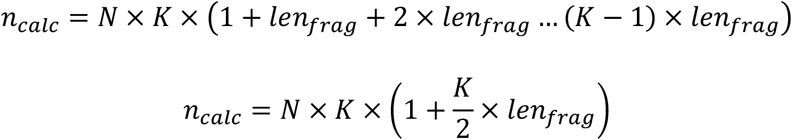

Where *len*_*frag*_ is the maximum length of the fragment.

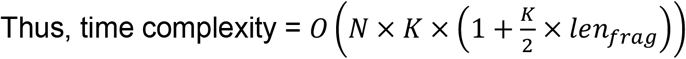

### 2.8 Predicted efficiency and fidelity values

For a set of overhangs called *Overhangs*, the net efficiency is computed as

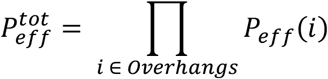

Predicted efficiency in percentage is computed by multiplying 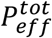 with 100. This represents the percentage efficiency w. r. t. the best overhang identified by Potapov et al.

Similarly, the net fidelity (probability of assembly without mismatch) is computed as

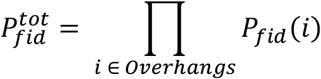

Predicted fidelity in percentage is computed by multiplying 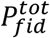 with 100. This represents the percentage of the population of the correct molecule from the set of all assembled molecules.

## 3 Results

### 3.1 Example

Consider a DNA sequence of length 35. The aim is to make K fragments between the range 10-13 and thus, in this case, *K* = 4. The fragments are indexed with *i* and DNA positions that form overhangs are indexed with *j*. The calculation starts at *i* = 0. Here *S(0,0)* is set to 1 to denote the starting site. Note that multiple start sites can be set by adding probabilities to different *j* positions.

At *i* = 1, only *j* in the range 10-13 are evaluated (*f*(*j*^*i*−1^) from section 2.2.2) (dotted grey lines in figure 1) as the rest of the positions do not qualify the fragment length criterion. The *j*^*i*−1^ which gives the highest value is picked as the trace (solid black and red arrows in Figure 1). For *i* = 2, the range of *j* evaluated is 20-26 (size range multiplied by *i*) as this range describes the shortest and largest fragments possible in relation to *i-1*. Similarly, the evaluation proceeds as per section 2.2.2 until *i > K* (section 2.2.3).

**Figure 1.**
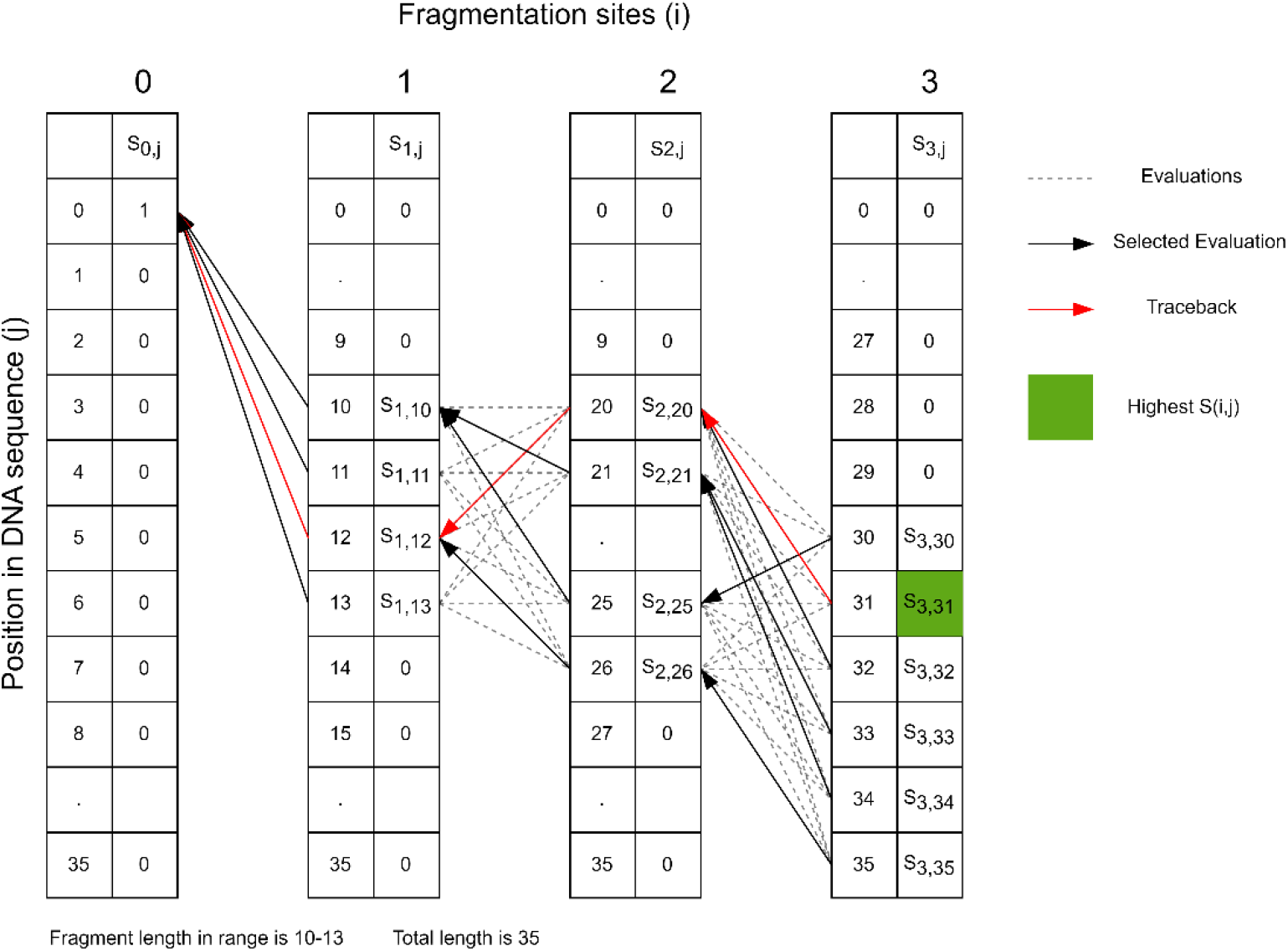
Illustration of OOGGA algorithm processing an example DNA sequence of length 35 for fragments of length range 10-13.

The traceback starts by evaluating all *i,j* positions that fall within the allowed fragment range. Assume that *S(3,31)* has the highest score in the example. The traceback follows the traces set during the evaluation stage starting from (3,31) until *i* = 0 (in this example (0,0) (solid red arrows in figure 1). The positions of overhangs identified are 10, 20 and 31 creating 4 fragments. Please note that the highest *S(i,j)* may not have the highest *i*. Thus, the algorithm can predict the optimal combination of overhangs and the number of fragments given only the DNA sequence and the size range of fragments.

### 3.2 Comparison of the fidelities of overhangs predicted by OOGGA and SplitSet

We compared the fidelity of overhangs from the fragments predicted by OOGGA and SplitSet. To do this we used three gene sequences 1) retinol binding protein 2) protein lp_0118 from *Lactobacillus plantarum* and 3) desmoplakin. The DNA sequences of these genes can be found in the supplementary data. SplitSet results were obtained from the web server after setting the required number of fragments as 5 (all other parameters were kept at their default set values). OOGGA results were obtained with *W*_*eff*_ set as 0 to optimize only the fidelity, and thus have a direct comparison with SplitSet. The number of fragments was set at 5, and the maximum and minimum fragment length was set as 20-150 for sequence 1, 20-100 for sequence 2 and 2000-3000 for sequence 3. The efficiency and fidelity values were calculated as shown in the methods section ‘Predicted efficiency and fidelity values’.

OOGGA predicts fragments with overhangs with higher or matching fidelities compared to SplitSet (Table 1) in all scenarios. This demonstrates the advantage of dynamic programming optimization by OOGGA, which guarantees the best solution over the stochastic search of SplitSet. However, since neither method optimize for the efficiency of ligation of the overhang, the efficiencies of the predicted overhangs predicted by both methods are poor (Table 2).

**Table 1.**
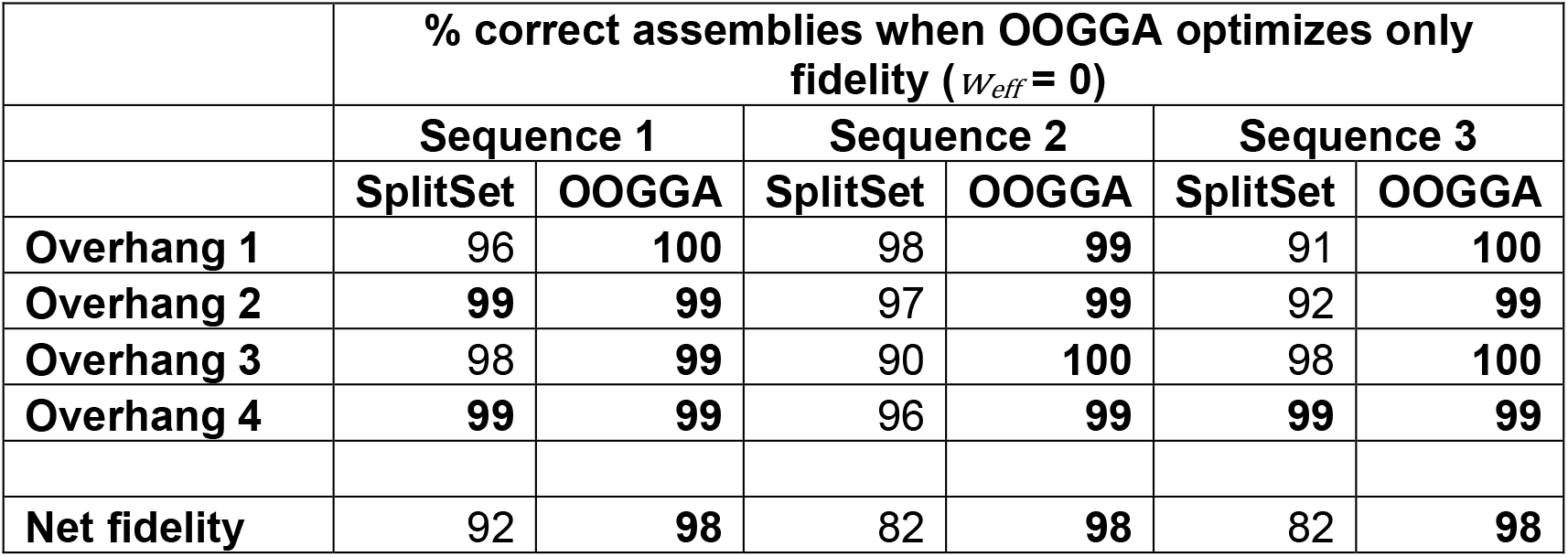
The predicted fidelity of overhangs predicted by SplitSet compared to OOGGA for the 3 DNA sequences. Fidelity is represented as % of correct assemblies. *W*_*eff*_ *= 0 and W*_*fid*_ *=* 1 for OOGGA. SplitSet values were obtained from its web server. Bold values show higher/matching values.

**Table 2.** The predicted efficiency of overhangs predicted by SplitSet compared to OOGGA for three DNA sequences. Efficiency is represented in % w. r. t. the most efficient overhang. *W*_*eff*_ *= 0 and W*_*fid*_ *=* 1 for OOGGA. SplitSet values were obtained from its web server. Bold values show higher/matching values.

| | % Efficiency w.r.t. most efficient overhang when OOGGA optimizes only fidelity ($w_{eff} = 0$) | | | | | |
| --- | --- | --- | --- | --- | --- | --- |
|  | Sequence 1 |  | Sequence 2 |  | Sequence 3 |  |
|  | SplitSet | OOGGA | SplitSet | OOGGA | SplitSet | OOGGA |
| Overhang 1 | **77** | 70 | 31 | **82** | **81** | 70 |
| Overhang 2 | **89** | 65 | **81** | 65 | **81** | 65 |
| **Overhang 3** | **85** | 66 | 57 | **70** | **85** | 30 |
| **Overhang 4** | 38 | **62** | **77** | 65 | **82** | 61 |
| **Net Efficiency** | **22** | 18 | 11 | **24** | **46** | 8 |

To check if OOGGA can predict overhangs that have both high efficiency and fidelity we set *W*_*eff*_ as 1.

### 3.3 OOGGA predicts overhangs with high fidelity and efficiency

We predicted overhangs with OOGGA for the three DNA sequences while setting *W*_*eff*_ as 1 and *W*_*fid*_ as 1 (equal preference to both efficiency and fidelity). With these weights, OOGGA Should optimize for overhangs with both high fidelity and efficiency. Other parameters used were the same as in section 3.2. SplitSet data used is the same as in section 3.2.

When *W*_*eff*_ was set as 1, OOGGA outperformed SplitSet in all efficiency-related scenarios (Table 3). However, this is expected as SplitSet does not optimize for efficiency. On evaluating the fidelity of the predicted overhangs, OOGGA predicts better overhangs for sequences 2 and 3 but falls behind SplitSet for sequence 1 (Table 4). However, it should be noted that the overhangs that OOGGA predicts are optimized for both efficiency and fidelity, compared to SplitSet which only optimizes for fidelity.

**Table 3.** The predicted efficiency of overhangs predicted by SplitSet compared to OOGGA for the 3 DNA sequences. Efficiency is represented in % w. r. t. the most efficient overhang. *W*_*eff*_ *= W*_*fid*_ *=* 1 for OOGGA. SplitSet values were obtained from its web server. Bold values show higher/matching values.

|  | **% Efficiency w.r.t. most efficient overhang when OOGGA optimizes fidelity and efficiency ($w_{eff} = 1, w_{fid} = 1$)** |  |  |  |  |  |
| --- | --- | --- | --- | --- | --- | --- |
|  | **Sequence 1** |  | **Sequence 2** |  | **Sequence 3** |  |
|  | **SplitSet** | **OOGGA** | **SplitSet** | **OOGGA** | **SplitSet** | **OOGGA** |
| **Overhang 1** | 77 | **95** | 31 | **90** | 81 | **100** |
| **Overhang 2** | 89 | **100** | 81 | **99** | 81 | **95** |
| **Overhang 3** | 85 | **90** | 57 | **100** | 85 | **93** |
| **Overhang 4** | 38 | **99** | 77 | **94** | 82 | **92** |
| **Net efficiency** | 22 | **84** | 11 | **85** | 46 | **82** |

**Table 4.** The predicted fidelity of overhangs predicted by SplitSet compared to OOGGA for the 3 DNA sequences. Fidelity is represented as % of correct assemblies. *W*_*eff*_ *= W*_*fid*_ *=* 1 for OOGGA. Splitset values were obtained from its web server. Bold values show higher/matching values.

|  | **% correct assemblies when OOGGA optimizes fidelity and efficiency ($w_{eff} = 1, w_{fid} = 1$)** |  |  |  |  |  |
| --- | --- | --- | --- | --- | --- | --- |
|  | **Sequence 1** |  | **Sequence 2** |  | **Sequence 3** |  |
|  | **SplitSet** | **OOGGA** | **SplitSet** | **OOGGA** | **SplitSet** | **OOGGA** |
| **Overhang 1** | **96** | 95 | 98 | **99** | 91 | **95** |
| **Overhang 2** | **99** | 95 | **97** | 93 | 92 | **95** |
| **Overhang 3** | **98** | 96 | 90 | **95** | 98 | **99** |
| **Overhang 4** | **99** | 93 | **96** | 95 | **99** | 98 |
| **Net fidelity** | **92** | 80 | 82 | **83** | 82 | **88** |

Thus, OOGGA predicts high-efficiency overhangs with fidelities comparable to SplitSet. OOGGA also allows use of various weights to bias efficiency and fidelity levels of the predictions. Such an example with *W*_*eff*_ *= 0*.*5 and W*_*fid*_ *=* 1 is show in appendix.

## Discussion

We have created a dynamic programming algorithm that predicts fragments with optimal overhangs for Golden Gate assembly (OOGGA). We compared it against NEBridge SplitSet^®^ which is the state-of-the-art method of predicting fragments for Golden gate assembly.

OOGGA outperforms or matches SplitSet in all scenarios on a direct comparison where both programs only optimize for fidelity. This is expected because OOGGA is a dynamic programming optimization algorithm while SplitSet is a stochastic search method, and the first guarantees the optimal solution. Moreover, OOGGA can predict overhangs with comparable accuracies to SplitSet even while optimizing for efficiency of ligation of overhangs (SplitSet does not optimize efficiency).

The difference in fidelity between OOGGA and SplitSet for each fragment is small enough that sequence upstream of the cleavage site can affect the outcome. However, on an average, the errors caused by the effect of the upstream site is equal on both OOGGA fragments and SplitSet fragments as they both used the same experimental data (Potapov, Jennifer L Ong, et al. 2018; Potapov, Jennifer L. Ong, et al. 2018) to predict fidelity. Since OOGGA is a dynamic programing predictor, it mathematically assures the best outcome (global minima) based on the provided data, which is not true for SplitSet. Thus, predictions from OOGGA will theoretically always (even if the difference is marginal) be better than SplitSet.

One might ask whether there is a need for such rigorous optimization. The following two paragraphs answer this question with answers distinct from academic curiosity.

Overhangs with high ligation efficiency would presumably reduce incubation times by an insignificant amount for day-to-day cloning. However, it is an important metric while designing overhangs for library preparations where higher efficiency implies a larger sample of the population. Thus, the overhangs generated by SplitSet are unsuitable for applications in large library preparations and OOGGA must be used for these applications as it predicts high efficiency and high-fidelity overhangs.

It is common to use fragments generated for Golden Gate assembly as part of modular systems. While optimizing for fidelity SplitSet only considers the mismatches between the predicted overhangs from the same sequence. Thus, these overhangs can have high error rates when combined with another system of fragments with a different set of overhangs. OOGGA avoids this by computing its fidelity score as the probability of an overhang of binding to only its Watson Crick match, mismatches with all overhangs are considered in doing this. Thus, the fidelity of the overhangs predicted by OOGGA is more universal compared to those predicted by SplitSet. Thus, overhangs of the fragments predicted by OOGGA are better in both the accuracy and efficiency of assembly.

These advantages of OOGGA can be experimentally verified by assembling two sets of fragments of a plasmid predicted by OOGGA and SplitSet separately, followed by counting the number of colonies after transformation (metric of efficiency) and selection by the corresponding antibiotic. Sequencing the plasmids isolated from the colonies can provide the rate of erroneous assembly (metric of fidelity). We are currently pursuing such an experiment.

OOGGA also supports the processing of DNA sequences with non-ATGC bases (eg. degeneracy codes), though the overhangs containing these bases are not considered for output. There can also be the exclusion of specific positions near these bases so that the overhangs are not predicted near them.

Note that the time complexity (section 2.3) and memory requirements of OOGGA, being a dynamic programming approach will be inferior compared to that of the SplitSet Monte Carlo method. However, designing fragments for Golden Gate assembly is not a time-sensitive event. The fidelity and efficiency of the fragments take priority over the increase in compute time and memory requirements.

Our results show that OOGGA is a better method compared to SplitSet to generate fragments for Golden Gate assembly. Our method predicts more accurate overhangs compared to SplitSet while also optimizing for efficiency and thus should be used for predicting overhangs for Golden Gate assembly instead of SplitSet.

## Acknowledgments

We would like to thank Dr. Raghavan Varadarajan and Dr. Nishad Matange for their critical comments of the work. SM would like to acknowledge CSIR SRF fellowship. This work was supported by the Department of Biotechnology, Government of India grant to the Indian Institute of Science Education and Research Pune; Department of Biotechnology, India under BIC grant (BT/PR40262/BTIS/137/38/2022).

## Competing interests

Neither SM nor MSM have any competing interests.

## Data availability

https://github.com/bigbigdumdum/OOGGA

## Appendix

Overhangs found by SplitSet vs OOGGA

Overhangs by SplitSet

- 1CBS: CTGA, AACC, AATG, AGAA
- 3K2Y: TTGT, GGTA, TAAG, CTGA
- 1LM7: TACC, GAAG, AATG, CGAA

Overhangs by OOGGA (*W*_*eff*_ = 0)

- 1CBS: AACA, GAGA, AGGA, CCAA
- 3K2Y: CGAA, GAGA, AACA, GCAA
- 1LM7: AACA, GAGA, ACAA, AGTA

Overhangs by OOGGA (*W*_*eff*_ = 1)

- 1CBS: ATCG, GATC, AGCC, CATG
- 3K2Y: CAAC, CATG, GATC, ATCG
- 1LM7: GATC, ATCG, AAAC, CGGA

Overhangs by OOGGA (*W*_*eff*_ = 0.5)

- 1CBS: CGGA, GATC, AGCC, CAAC
- 3K2Y: CAAC, CATG, GATC, ATCG
- 1LM7: AAAC, CGGA, GATC, ATCC

**SplitSet vs OOGGA at *W*_*eff*_ = 0.5**

**Table 5.** The predicted efficiency of overhangs predicted by SplitSet compared to OOGGA for the 3 DNA sequences. Efficiency is represented in % w. r. t. the most efficient overhang. *W*_*eff*_ *= 0*.*5 and W*_*fid*_ *=* 1 for OOGGA. Splitset values were obtained from its web server. Bold values show higher/matching values.

| % Efficiency w.r.t. most efficient overhang ($w_{eff} = 0.5$) | | | | | | |
| --- | --- | --- | --- | --- | --- | --- |
|  | Sequence 1 |  | Sequence 2 |  | Sequence 3 |  |
|  | SplitSet | OOGGA | SplitSet | OOGGA | SplitSet | OOGGA |
| Overhang 1 | 77 | **92** | 31 | **90** | 81 | **93** |
| Overhang 2 | 89 | **100** | 81 | **99** | 81 | **92** |
| Overhang 3 | **85** | 83 | 57 | **100** | 85 | **100** |
| Overhang 4 | 38 | **90** | 77 | **95** | 82 | **90** |
| Net Efficiency | 22 | **74** | 11 | **85** | 46 | **77** |

**Table 6.** The predicted fidelity of overhangs predicted by SplitSet compared to OOGGA for the 3 DNA sequences. Fidelity is represented as % of correct assemblies. *W*_*eff*_ *= 0*.*5 and W*_*fid*_ *=* 1 for OOGGA. Splitset values were obtained from its web server. Bold values show higher/matching values.

| % correct assemblies ($w_{eff} = 0.5$) | | | | | | |
| --- | --- | --- | --- | --- | --- | --- |
|  | Sequence 1 |  | Sequence 2 |  | Sequence 3 |  |
|  | SplitSet | OOGGA | SplitSet | OOGGA | SplitSet | OOGGA |
| Overhang 1 | 96 | **98** | 98 | **99** | 91 | **99** |
| Overhang 2 | **99** | 95 | **97** | 93 | 92 | **98** |
| **Overhang 3** | **98** | 96 | 90 | **95** | **98** | 95 |
| **Overhang 4** | **99** | **99** | **96** | 95 | **99** | **99** |
| **Net fidelity** | **92** | 88 | 82 | **83** | 82 | **91** |

## Notes

https://github.com/bigbigdumdum/OOGGA

### Competing Interest Statement

The authors have declared no competing interest.

## Reference

Bilotti, Katharina, Vladimir Potapov, John M. Pryor, Alexander T. Duckworth, James L. Keck, and Gregory J. S. Lohman. 2022. “Mismatch Discrimination and Sequence Bias during End-Joining by DNA Ligases.” Nucleic Acids Research 50(8):4647–58. doi: 10.1093/nar/gkac241.

Engler, Carola, Romy Kandzia, and Sylvestre Marillonnet. 2008. “A One Pot, One Step, Precision Cloning Method with High Throughput Capability.” PLOS ONE 3(11):e3647.

Potapov, Vladimir, Jennifer L. Ong, Rebecca B. Kucera, Bradley W. Langhorst, Katharina Bilotti, John M. Pryor, Eric J. Cantor, Barry Canton, Thomas F. Knight, Thomas C. Evans, and Gregory J. S. Lohman. 2018. “Comprehensive Profiling of Four Base Overhang Ligation Fidelity by T4 DNA Ligase and Application to DNA Assembly.” ACS Synthetic Biology 7(11):2665–74. doi: 10.1021/acssynbio.8b00333.

Potapov, Vladimir, Jennifer L Ong, Bradley W. Langhorst, Katharina Bilotti, Dan Cahoon, Barry Canton, Thomas F. Knight, Thomas C. Evans Jr, and Gregory J. S. Lohman. 2018. “A Single-Molecule Sequencing Assay for the Comprehensive Profiling of T4 DNA Ligase Fidelity and Bias during DNA End-Joining.” Nucleic Acids Research 46(13):e79–e79. doi: 10.1093/nar/gky303.

Pryor, John M., Vladimir Potapov, Katharina Bilotti, Nilisha Pokhrel, and Gregory J. S. Lohman. 2022. “Rapid 40 Kb Genome Construction from 52 Parts through Data-Optimized Assembly Design.” ACS Synthetic Biology 11(6):2036–42. doi: 10.1021/acssynbio.1c00525.

Pryor, John M., Vladimir Potapov, Rebecca B. Kucera, Katharina Bilotti, Eric J. Cantor, and Gregory J. S. Lohman. 2020. “Enabling One-Pot Golden Gate Assemblies of Unprecedented Complexity Using Data-Optimized Assembly Design.” PLOS ONE 15(9):e0238592.

